# Microbial metabolite fluxes in a model marine anoxic ecosystem

**DOI:** 10.1101/625087

**Authors:** Stilianos Louca, Yrene M. Astor, Michael Doebeli, Gordon T. Taylor, Mary I. Scranton

## Abstract

Permanently anoxic regions in the ocean are widespread, and exhibit unique microbial metabolic activity exerting substantial influence on global elemental cycles and climate. Reconstructing microbial metabolic activity rates in these regions has been challenging, due to the technical difficulty of direct rate measurements. In Cariaco Basin, which is the largest permanently anoxic marine basin and an important model system for geobiology, long-term monitoring has yielded time series for the concentrations of biologically important compounds; however the underlying metabolite fluxes remain poorly quantified. Here we present a computational approach for reconstructing vertical fluxes and *in situ* net production/consumption rates from chemical concentration data, based on a 1-dimensional time-dependent diffusive transport model that includes adaptive penalization of overfitting. We use this approach to estimate spatiotemporally resolved fluxes of oxygen, nitrate, hydrogen sulfide, ammonium, methane and phosphate within the sub-euphotic Cariaco Basin water column (depths 150–900 m, years 2001–2014), and to identify hotspots of microbial chemolithotrophic activity. Predictions of the fitted models are in excellent agreement with the data, and substantially expand our knowledge of the geobiology in Cariaco Basin. In particular, we find that the diffusivity, and consequently fluxes of major reductants such as hydrogen sulfide and methane, are about two orders of magnitude greater than previously estimated, thus resolving a long standing apparent conundrum between electron donor fluxes and measured dark carbon assimilation rates.

## 1 Introduction

Permanently or temporarily anoxic regions in the ocean are a topic of increasing interest due to their unique microbial ecology (Wright *et al.*, 2012; Ulloa *et al.*, 2013), their importance to global elemental cycles and marine productivity (Ulloa *et al.*, 2012), and the intensifying deoxygenation of the ocean (Schmidtko *et al.*, 2017; Breitburg *et al.*, 2018). Microorganisms in these regions are adapted to operate under oxygen-limited or oxygen-depleted conditions, making use of alternative terminal electron acceptors for respiration and often utilizing inorganic substrates for energy. In many anoxic marine zones, reductants such as hydrogen sulfide, ammonium and the potent greenhouse gas methane, diffusing upwards from underlying layers or the sediments, react biologically with oxidants such as oxygen and nitrate produced in the overlying layers, thus fueling chemolithoautotrophic activity and affecting marine nitrogen, sulfur, oxygen and carbon budgets (Ulloa *et al.*, 2012; Taylor *et al.*, 2018). In the Cariaco Basin, a permanently anoxic marine region off the coast of Venezuela, multi-decadal monitoring has generated rich time series of the distribution of metabolically important compounds over space and time (Scranton *et al.*, 2014; Muller-Karger *et al.*, 2019). These data revealed the existence of a strong dynamic redox gradient over depth, along which upward diffusing reductants such as hydrogen sulfide are directly or indirectly oxidized by oxidants such as oxygen in a transition zone roughly spanning depths 200-400 m, sometimes referred to as “redoxcline” (Ho *et al.*, 2004; Li *et al.*, 2012b; Taylor *et al.*, 2018). Concurrent molecular surveys revealed unique microbial communities that exhibit a clear spatial organization across depth, and elevated population densities within the redoxcline (Taylor *et al.*, 2006; Rodriguez-Mora *et al.*, 2015; Taylor *et al.*, 2018; Cernadas-Martín *et al.*, 2017; Suter *et al.*, 2018). However, chemical fluxes across space and microbial metabolic rates in Cariaco Basin and other anoxic regions remain poorly quantified and are largely temporally unresolved, thus making a mechanistic connection between chemical transitions and microbial ecological dynamics difficult (Taylor *et al.*, 2018). A major reason for this gap in our knowledge is that, compared to chemical concentration measurements, explicit metabolic rate measurements are technically challenging, especially when performed *in situ*.

Mathematical modeling is sometimes used to indirectly estimate the flux rates that underly the observed chemical concentrations (Scranton *et al.*, 1987; Berg *et al.*, 1998; Taylor *et al.*, 2001; Samodurov *et al.*, 2013; Li *et al.*, 2012b; Cernadas-Martín *et al.*, 2017; Taylor *et al.*, 2018). For example, Scranton *et al.* (1987) used a time-dependent diffusion box model to estimate sulfide fluxes from the sediments into the Cariaco Basin water column. These calculations, however, were based solely on measurements at two time points (in 1973 and 1982), ignored possible *in situ* sulfide production (Li *et al.*, 2012b), and assumed that sulfide was consumed entirely at some fixed depth, thus ignoring shifts in the redoxcline depth over time (Scranton *et al.*, 2014). On the other hand, Li *et al.* (2012b) and Cernadas-Martín *et al.* (2017) used a 1-dimensional diffusion model for Cariaco Basin to estimate fluxes of various compounds produced or consumed during microbial metabolism (henceforth “metabolites” for simplicity). Their models only estimated fluxes into and out of a narrow depth interval (~100–200 m wide), thus missing possible metabolic activity at other depths, and assumed that metabolite depth profiles were at steady state, thus ignoring possible temporal lags in the response of redox gradients to flux changes (Scranton *et al.*, 1987, 2014) and microbial dynamics (Taylor *et al.*, 2018).

Here we develop a computational approach for estimating metabolite fluxes and net production/consumption rates across space and time, using chemical concentration data measured at arbitrary spacetime points. Our approach is based on a 1-dimensional time- and depth-dependent diffusive transport model that accounts for temporal changes in boundary conditions, diffusive transport coefficients and *in situ* production rates, as well as for potential geometric dilution effects due to variation of a system’s lateral (cross-sectional) area with depth. We use our approach to reconstruct spatiotemporally resolved metabolite fluxes across the sub-euphotic Cariaco Basin water column (depths 150–900 m) during the years 2001–2014. We consider several important metabolites, including oxygen (O_2_), nitrate 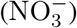, hydrogen sulfide (H_2_S), ammonium 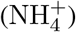, methane (CH_4_) and phosphate 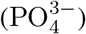. Our estimates yield detailed insight into the microbial activity that underlies the geochemical structure of the Cariaco Basin water column.

## 2 Results and Discussion

### 2.1 Estimating diffusivity over space and time

In largely stagnant marine basins such as Cariaco Basin (Scranton *et al.*, 1987; Samodurov *et al.*, 2013; Taylor *et al.*, 2018), the Black Sea (Ivanov and Samodurov, 2001) and parts of the Arabian Sea (Lam *et al.*, 2011), eddy (turbulent) diffusion is the dominant mode of vertical transport of dissolved metabolites. Great uncertainty often exists over the magnitude of the vertical diffusion coefficient (henceforth “diffusivity”), and this uncertainty can substantially influence flux estimates. Previous theoretical and empirical work suggests that the diffusivity (denoted *D*) in such water columns is typically related to the buoyancy frequency (denoted *N*) through a power-law relationship of the form:

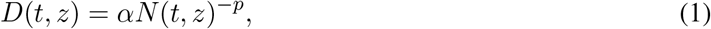

where *t* is time, *z* is depth, and *α* and *p* are system-specific parameters (Sarmiento *et al.*, 1976; Gregg, 1977; Armi, 1979; Smethie, 1980; Osborn, 1980; Gargett and Holloway, 1984; Gargett, 1984; Gregg *et al.*, 1986). Such a power law can be mathematically justified for stably stratified systems without double diffusion and in which the bulk of kinetic energy reaches turbulent scales via internal waves (Gargett, 1984). In these scenarios, the parameter *α* accounts for the average energy entering the system (e.g., by winds or tides), and *p* (typically between 0.5 and 1) reflects the a priori unknown *N*-dependence of internal wave velocity variances (Gargett, 1984). Alternatively, a power law can be derived for stably stratified systems in which an apparent diapycnal diffusion-like mixing is caused mainly by turbulence near the basin bottom/boundaries and topographic features and a rapid redistribution of material throughout the interior by isopycnal advection (Armi, 1979), yielding *p* ≈ 2. In some studies, a power law relationship between *D* and *N* is a purely empirical observation, with *p* ranging between 0.5 and 2 (Sarmiento *et al.*, 1976; Svensson, 1980; Gargett, 1984). We mention that alternative *N*-dependent models for *D* have also been derived, based on different assumptions regarding the origin and dissipative nature of kinetic energy (Munk and Anderson, 1948; Lee *et al.*, 2006; Olbers and Eden, 2013).

Equation (1) has been used extensively to predict turbulent transport of dissolved gases and salts in various systems, especially in anoxic marine systems (Fennel and Boss, 2003; Ho *et al.*, 2004; Li *et al.*, 2012b; Samodurov *et al.*, 2013; Reed *et al.*, 2014; Louca *et al.*, 2016). The buoyancy frequency *N* can be calculated from the measured salinity and temperature profiles, however the appropriate values for *α* and *p* are typically poorly constrained. Some studies have estimated the parameters *α* and *p*, or diffusivity itself, using spatiotemporal profiles of salinity (Gade, 1970; Smethie, 1980; Lewis and Perkin, 1982; Ivanov and Samodurov, 2001) or other conserved tracers (Svensson, 1980; Gargett, 1984). For example, Svensson (1980) used the diffusion of Rhodamine B as a semi-conserved tracer to estimate a power-law exponent of *p* = 1.2 in Byfjorden (Sweden), Gade (1970) used the salt budget to estimate an exponent of *p* = 1.6 in Oslofjord (Norway), and Lewis and Perkin (1982) used the salt budget to estimate *D* in Agfardlikavsa Fjord (Greenland), revealing a power law dependence on *N* with an exponent *p* ≈ 1.2 (Gargett, 1984). In Cariaco Basin analogous parameter estimates are lacking, and previous studies simply assumed an exponent of *p* = 1 (Scranton *et al.*, 1987; Taylor *et al.*, 2001; Ho *et al.*, 2004; Li *et al.*, 2012b; Samodurov *et al.*, 2013; Taylor *et al.*, 2018). A value of *p* = 1 is also frequently assumed in other systems (Zopfi *et al.*, 2001; Yakushev *et al.*, 2007; Yakushev, 2013), although some studies instead assumed *p* = 2 (Fennel and Boss, 2003; Reed *et al.*, 2014). The factor *α* is usually chosen roughly based on estimates from other marine systems (Ho *et al.*, 2004; Li *et al.*, 2012b; Samodurov *et al.*, 2013; Taylor *et al.*, 2018).

Here, to estimate both *α* and *p* for Cariaco Basin, thus resolving a major source of uncertainty in flux estimates, we used a 1-dimensional diffusive transport model for the salt budget in Cariaco Basin, and fitted the parameters *α* and *p* by minimizing the deviation of the model predictions from salinity measurements. Specifically, for any given choice of *α* and *p*, we numerically solved the diffusion equation

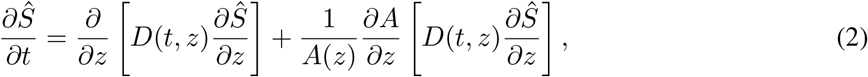

where *Ŝ* is the predicted salinity, *D* is the diffusivity calculated using Eq. (1) and *A*(*z*) is the lateral (cross-sectional) area of the basin at depth *z* (Supplemental Fig. S.1). The last term in Eq. (2) accounts for geometric dilution effects due to variation of the basin area over depth (Samodurov *et al.*, 2013). This model omits occasional lateral intrusions of Caribbean Sea water (Samodurov *et al.*, 2013; Scranton *et al.*, 2014); the accuracy of the model is assessed in retrospect. We considered the depth range 150–900 m and the period spanning years 2001–2014, with boundary conditions provided by measured salinities at 150 m and 900 m. This depth range was chosen because 150 m is the maximum depth of the sill separating Cariaco Basin from the open ocean (and above which non-diffusive salt transport due to lateral currents is more pronounced), and because 900 m is the depth of the saddle that separates the west and east sub-basins in Cariaco (Taylor *et al.*, 2001). The parameters *α* and *p* were gradually adjusted using an optimization algorithm, so that the sum of squared deviations between *Ŝ* and the measured salinity is minimized. This yielded the estimates *α* = 0.0001316 and *p* = 1.7433, when *D* is measured in cm^2^ · s^−1^ and *N* is measured in s^−1^. The agreement between the predicted and measured salinity profiles was excellent, as measured by the fraction of explained variance (*r*^2^=0.982, Supplemental Fig. S.3B). This suggests that neglected processes, such as occasional lateral water intrusions, only have a minor influence on the Cariaco Basin salt budget during the considered time interval.

To test whether our diffusivity estimates are sensitive to the choice of model, we also considered an alternative model known as Munk-Anderson scheme (Munk and Anderson, 1948; James, 1977; Lee *et al.*, 2006):

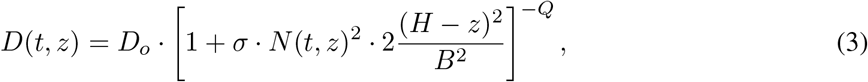

where *D*_*o*_, *σ*, *Q* and *B* are system-specific model parameters and *H* is the bottom depth (*H* ≈ 1400 m for Cariaco Basin). The Munk-Anderson scheme assumes that diffusion-like vertical mixing is driven by frictional velocity shear, induced by tidal motions damped near the basin bottom (Munk and Anderson, 1948). The specific formula in Eq. (3) is based on an empirical logarithmic profile of horizontal current velocities (James, 1977) and was used by Lee *et al.* (2006) in a North Atlantic ocean model. When we fit the above model to the salinity profiles in Cariaco Basin (depths 150–900 m, years 2001–2014), we obtained very similar diffusivity estimates as with the power law model (Supplemental Figs. S.4A,B). Further, when we combined both models into a single additive model (i.e., using the sum of Eq. 1 and Eq. 3), we again obtained similar diffusivity estimates as before (Supplemental Figs. S.4C,D). While our findings do not resolve which model provides the most suitable mechanistic explanation of mixing in Cariaco Basin, all models yield similar estimates for the effective diffusivity.

To further confirm the robustness of our model-based estimates we also considered a model-independent approach, in which diffusivity is estimated directly from the salinity data (*S*) regardless of the buoyancy frequency and without assuming a particular process as the cause of mixing (Fig. 1C, details in Supplement S.1). Briefly, this alternative approach assumes that *D* is constant over time and somehow known at some given “anchor depth” *z*_*a*_. In this case, *D*(*z*) can be estimated using the implicit formula:

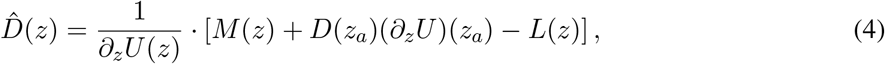

where *M*(*z*), *U*(*z*) and *L*(*z*) are auxiliary quantities calculated using the salinity data, defined as follows:

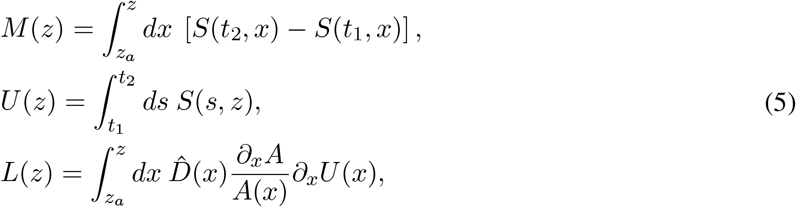

and where *t*_1_ < *t*_2_ are any two time points. The accuracy of the “anchored” estimate 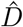 defined in Eq. (4) improves for larger considered time spans |*t*_2_ − *t*_1_|, and hence we used the full available time range (years 2001–2014). Because the auxiliary variable *L* itself depends on the estimated diffusivity 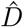, an iterative approach was used to solve for 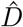. The choice of *z*_*a*_ is in principle arbitrary, as long as *D*(*z*_*a*_) can be determined somehow. A similar approach was used previously by Samodurov *et al.* (2013) to estimate *D* in Cariaco Basin, using the anchor depth *z*_*a*_ = 150 m, with an important difference: *Samodurov et al.* estimated *D*(*z*_*a*_) using the buoyancy-frequency-based formula in Eq. (1), with *α* based on other marine systems and assuming *p* = 1, whereas here we made no assumption about *D*(*z*_*a*_) and instead estimated *D*(*z*_*a*_) from the salinity data via least-squares fitting (details in Supplement S.1). We emphasize that this estimate is strictly speaking only valid if the true diffusivity *D* does not vary with time, and hence it should only be used as a rough sanity check. This alternatively estimated diffusivity profile again closely reproduces the measured salinity profile (*r*^2^ = 0.982, Supplemental Fig. S.3C) and also approximately resembles our previous diffusivity estimates (Fig. 1B), further increasing our confidence in these estimates.

**Figure 1:**
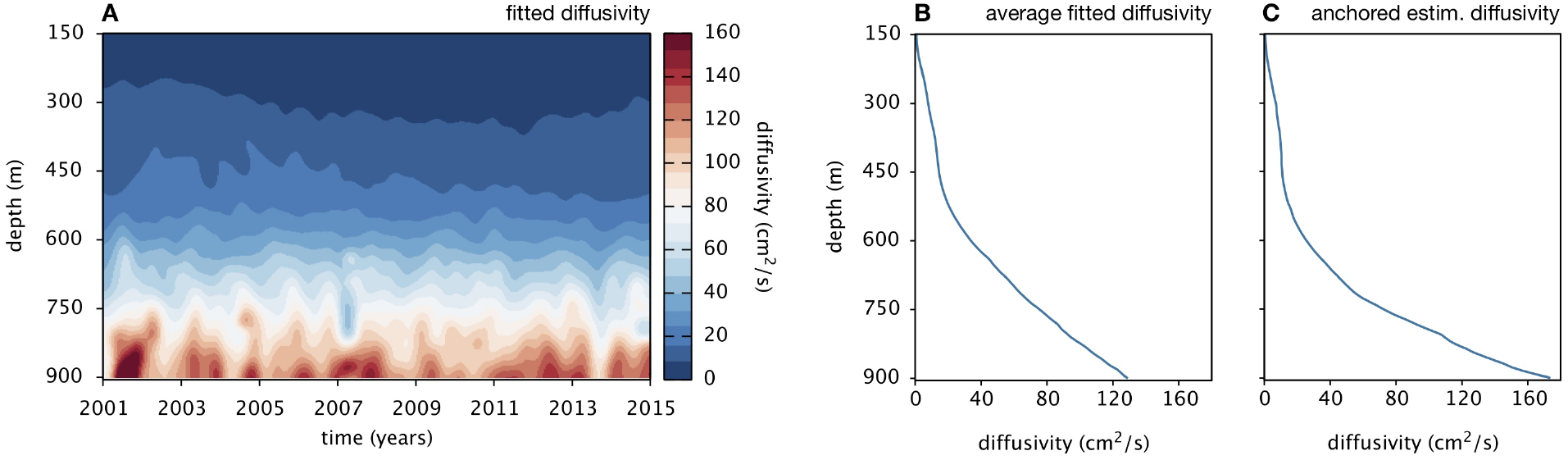
Estimated diffusivity in Cariaco Basin. (A) Diffusivity in Cariaco Basin (station CARIACO) over depth and time, estimated based on the buoyancy frequency, using the power law in Eq. (1) and the fitted parameters *α* = 0.0001316 and *p* = 1.7433. (B) Time-averaged diffusivity depth profile, calculated from A. (C) Estimated diffusivity depth-profile, estimated using Eq. (4) and assuming that *D* is independent of time.

As seen in Fig. 1A, the estimated diffusivity increases drastically with depth, due to the decreased buoyancy frequency and the super-linear scaling of *D* (*p* > 1). Our diffusivity estimates, particularly those near the bottom, are substantially higher than typical diffusivities estimated in other fjords and basins (Gargett, 1984; Yakushev *et al.*, 2007). In Cariaco Basin, stratification is extremely weak towards the bottom, allowing for more rapid diapycnal mixing than in the redoxcline. An exponent *p* greater than 1 and closer to 2 also suggests that mixing in Cariaco Basin is largely driven by turbulence near the basin’s boundaries (Armi, 1979), especially at depth where the lateral area decreases substantially (Supplemental Fig. S.1). Compared to other prominent anoxic marine systems, Cariaco Basin is relatively compact, with a horizontal area (at the sill’s depth) about 40 times smaller than the Black Sea (Kideys, 2002) and the Baltic Sea (Leppäranta and Myrberg, 2009) and about 130 times smaller than the Arabian Sea (Goyet *et al.*, 1998), potentially resulting in stronger boundary mixing than in those other systems. We point out that, in reality, the eddy diffusivity may exhibit substantial lateral heterogeneity, and may be greater near the Basin’s walls than in the center. The diffusivity profile estimated here thus represents the effective (laterally averaged) diffusivity under a 1-dimensional transport model that describes laterally-averaged vertical fluxes. Such a model is itself only valid under the implicit assumption that lateral mixing is much faster than vertical mixing — a reasonable assumption for Cariaco Basin, since stratification is largely vertical.

Our diffusivity estimates are substantially higher than estimates from previous studies in Cariaco Basin, all of which assumed an exponent of *p* = 1 (Scranton *et al.*, 1987; Li *et al.*, 2012b; Samodurov *et al.*, 2013; Taylor *et al.*, 2018). An exponent greater than 1 (*p* ≈ 1.7) is strongly supported by our fitted model, as the value *p* = 1 results in a much lower goodness of fit (Supplemental Fig. S.13). An exponent *p* > 1 is also consistent with our alternatively estimated diffusivity profiles (Fig. 1C and Supplemental Fig. S.4). Hence, previous studies probably underestimated *D* in Cariaco Basin, especially in deeper waters. As we discuss below, this has substantial implications for metabolite flux estimates and may explain some of the apparent imbalances between metabolite supply and biological demand previously observed in Cariaco Basin.

### 2.2 Inverse linear transport modeling

Our approach for estimating vertical fluxes and volume-specific net production rates of dissolved metabolites is based on the following 1-dimensional reaction-diffusion differential equation for the metabolite’s volumetric concentration:

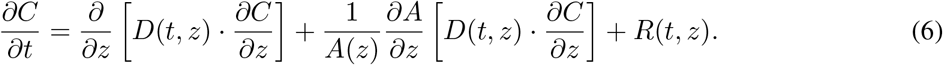

Here, *C*(*t, z*) is the metabolite’s concentration (mol·L^−1^), *D*(*t, z*) is the diffusivity (e.g., as estimated above), *A*(*z*) is the lateral (cross-sectional) area of the basin and *R*(*t, z*) is the a priori unknown volume-specific net metabolite production rate at any time *t* and depth *z*. Similarly to the salinity model above, this model can account for geometric dilution effects due to variations of the lateral basin area with depth (Samodurov *et al.*, 2013). Variants of the above model have been used extensively to describe dissolved nutrient transport in Cariaco Basin (Scranton *et al.*, 1987; Li *et al.*, 2012b; Cernadas-Martín *et al.*, 2017; Taylor *et al.*, 2018), although previous studies made simplifying assumptions such as that *R* was negligible (Scranton *et al.*, 1987), that *D* was constant over time (Li *et al.*, 2012b; Taylor *et al.*, 2018) or that *C* was at steady state (i.e., *∂C/∂t* = 0; Li *et al.*, 2012b; Cernadas-Martín *et al.*, 2017; Taylor *et al.*, 2018). Since *C* has been measured and *D* has been previously estimated, in principle one could directly calculate the unknown rate *R* at various times and depths through a simple algebraic reordering of Eq. (6). Unfortunately, this approach generally suffers from high estimation errors. The main reason is that the numerical estimation of spatial derivatives from discrete depth profiles, or of temporal derivatives from discrete time series, typically leads to an amplification of high-frequency noise (Knowles and Renka, 2014).

An alternative approach for estimating *R* that reduces estimation noise and avoids the risk of overfitting is to choose *R* on a finite spatiotemporal grid (“fitting grid”), such that the corresponding predicted distribution *Ĉ* obtained by solving the differential equation (6) best matches the observed profile *C*. This approach, known as “inverse linear transport modeling” (ILTM), is widely used in oceanography and atmospheric sciences, where known distributions of compounds are used to estimate unknown sources and sinks (Berg *et al.*, 1998; Houweling *et al.*, 1999; Mikaloff Fletcher *et al.*, 2006; Hirsch *et al.*, 2006; Mikaloff Fletcher *et al.*, 2007; Steinkamp, 2011; Lam *et al.*, 2011; Lettmann *et al.*, 2012; Martinez-Camara *et al.*, 2014; Louca *et al.*, 2016). We mention that most existing studies — including those investigating metabolite fluxes in anoxic water columns or sediments (Berg *et al.*, 1998; Lam *et al.*, 2011; Lettmann *et al.*, 2012; Louca *et al.*, 2016) — assumed that *C* was at steady state even when fluxes were estimated at multiple time points, however this assumption may be needlessly and overly restrictive. To reduce spurious oscillations in the estimated *R* (a common ILTM artifact), excessively high estimates of *R* that only marginally improve the agreement with the data are penalized, a procedure known as Tikhonov regularization (Björck, 1996; Hansen, 2000; Lettmann *et al.*, 2012). Specifically, for any given metabolite, the vector containing all values of *R* on the fitting grid (denoted **R**) is estimated by minimizing the expression:

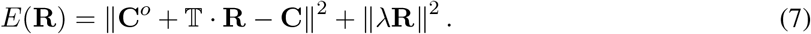

Here, **C** is a vector listing measured concentrations at arbitrary spacetime points, **C**^*o*^ is a pre-calculated vector listing concentrations predicted in the absence of any net production (i.e., when *R* = 0 and accounting for initial and boundary conditions, Supplemental Fig. S.7), ║·║^2^ denotes the squared norm of a vector (i.e., the sum of squares of all its components) and *E*(**R**) denotes the function to be minimized by appropriate choice of **R**. The matrix 𝕋 maps net production rates on the fitting grid to concentrations on the same spacetime points as the data, and is pre-calculated using the differential equation (6). The first ║·║^2^ term in Eq. (7) corresponds to the deviation of the predicted concentrations from the data, while the second ║·║^2^ term corresponds to the overall magnitude of the estimated net production rates. The “regularization factor” *λ* modulates the penalization of spurious rate estimates, balanced against achieving a better fit to the data, and is chosen adaptively and separately for each metabolite depending on the data using a cross-validation algorithm (Golub *et al.*, 1979). Hence, for a chosen *λ*, the task of estimating net production rates based on concentration data translates to an optimization problem, which can be solved numerically using linear algebra software (Supplement S.4). Because all data points **C** are used concurrently to fit the full spatiotemporal rate profile **R**, this method is more robust against measurement errors than previous methods that only use data from a single time point at a time (Berg *et al.*, 1998; Lam *et al.*, 2011; Lettmann *et al.*, 2012; Louca *et al.*, 2016).

We emphasize that the resolution and placement of the fitting grid must be chosen carefully to avoid the risk of overfitting. Indeed, the number of spacetime points on the fitting grid dictates the number of fitted free parameters, and hence the fitting grid must be much coarser than the concentration data **C**. At the same time, the fitting grid should capture the major variations in *R* over space and time, as indicated in the concentration data. Hence, the fitting grid should be densest in those spacetime regions where *R* is suspected to vary most and where, ideally, concentration measurements are also densest. The latter constraint underscores the importance of carefully choosing the times and depths targeted by oceanographic surveys. In practice, the fitting grid may need to be revised through trial and error and using expert knowledge of the system, for example where substantial oscillations in the estimated rates are obviously spurious. A common and useful difference between spurious and true variations in the estimated *R* is that the former tend to be much more sensitive to small variations in the fitting grid.

### 2.3 Metabolite fluxes in Cariaco Basin

We used the above ILTM approach to estimate net *in situ* production rates of several important dissolved metabolites in the Cariaco Basin sub-euphotic water column, using concentration time series spanning depths 150–900 m and years 2001–2014 (Fig. 2A–F). In addition to production rate estimates, we also estimated vertical metabolite flux rates from the bottom (depths>900 m) and from the overlying waters (depths<150 m) into the sub-euphotic zone and into the redoxcline. Estimated net metabolite production rates, interpolated between grid points, are shown in Fig. 3. The agreement between the measured metabolite concentrations and those predicted based on the estimated net production rates was generally good, with a fraction of explained variance (*r*^2^) between 0.878 and 0.973 depending on the metabolite (Fig. 2). The main features not captured by the fitted models are rapid fluctuations constrained within small depth intervals, potentially originating from occasional lateral water intrusions (Scranton *et al.*, 2014; Muller-Karger *et al.*, 2019), as well as seasonally driven variations in oxygen and nitrate concentrations (Fig. 2). In contrast, the fitted models accurately capture major decadal trends, most prominently seen in the sulfide and methane profiles (Figs. 2C,E). The inability of the fitted models to capture rapid transient small-scale fluctuations stems from two fundamentally information-theoretical limitations: First, the spatiotemporal resolution of the available data imposes a bound on the resolution of the fitting grid on which *R* can be independently estimated, and greater grid resolutions would substantially increase the risk of overfitting. Second, the estimation of *R* mathematically corresponds to an inversion (specifically, a deconvolution) of the diffusion process, and hence tends to amplify high-frequency noise in the data; the amplified noise manifests as spurious rapid oscillations in the estimated rates that are hard to distinguish from real fluctuations (Steinkamp, 2011; Lettmann *et al.*, 2012). The temporal resolution of our estimates is thus constrained by the time scales associated with diffusive mixing in Cariaco Basin which, based on the typical travel times of diffusing particles between the bottom boundary and the redoxcline, are in the order of ~2.4 years (Supplement S.2). Thus, estimated metabolic rates at any time point should be seen as local temporal averages over those time scales.

**Figure 2:**
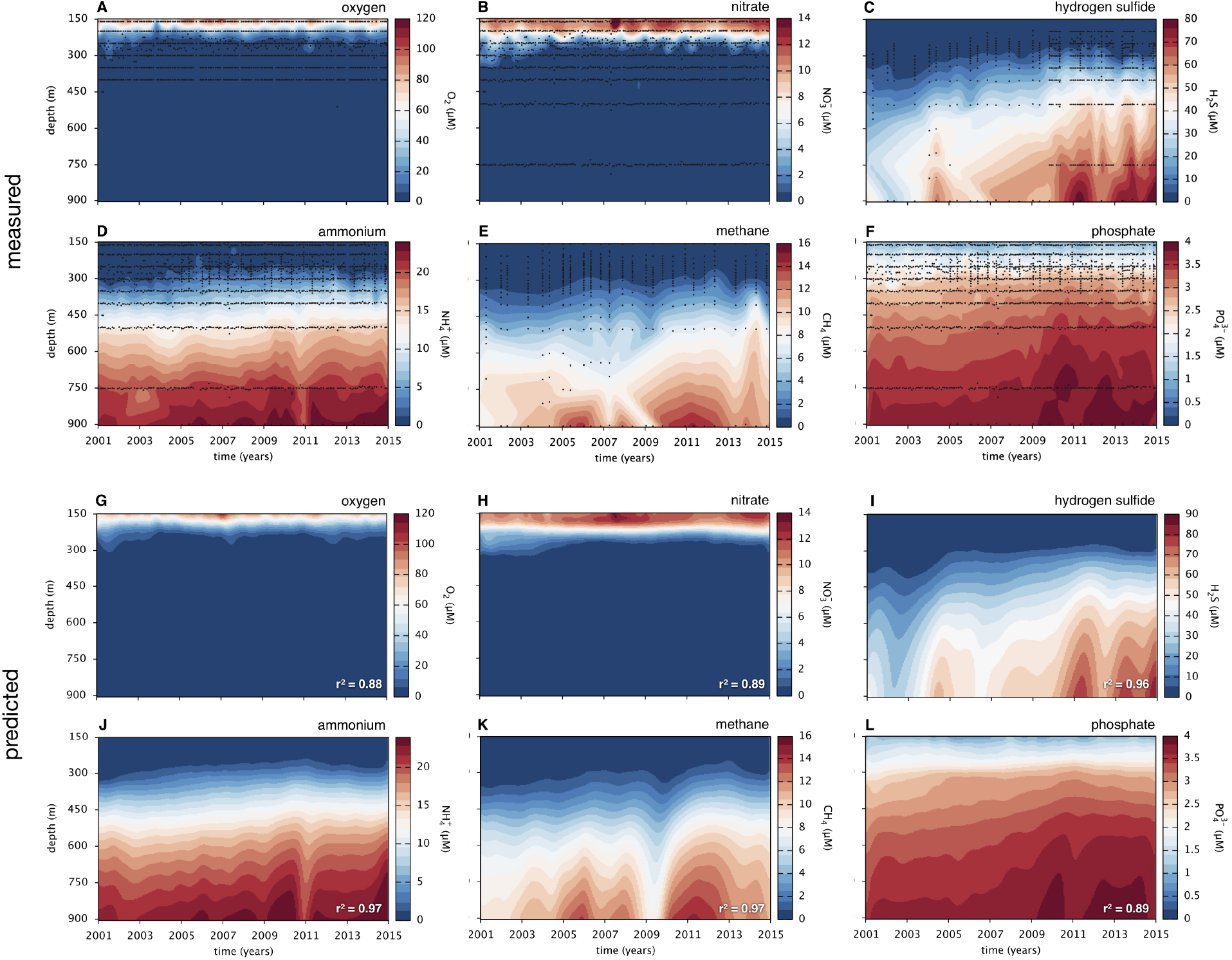
Metabolite concentrations in Cariaco Basin (data versus fitted models). A–F: Measured metabolite concentrations in Cariaco Basin (station CARIACO) over depth and time (A: oxygen, B:nitrate, C:hydrogen sulfide, D:ammonium, E:methane, F:phosphate). Black dots denote data points; contour plots are bilinear interpolations between data points. Data sources are described in the Methods. G–L: Predicted metabolite concentrations, based on the net production rates estimated via ILTM. Fractions of explained variance (*r*^*2*^), when compared to the data, are indicated in the figures.

**Figure 3:**
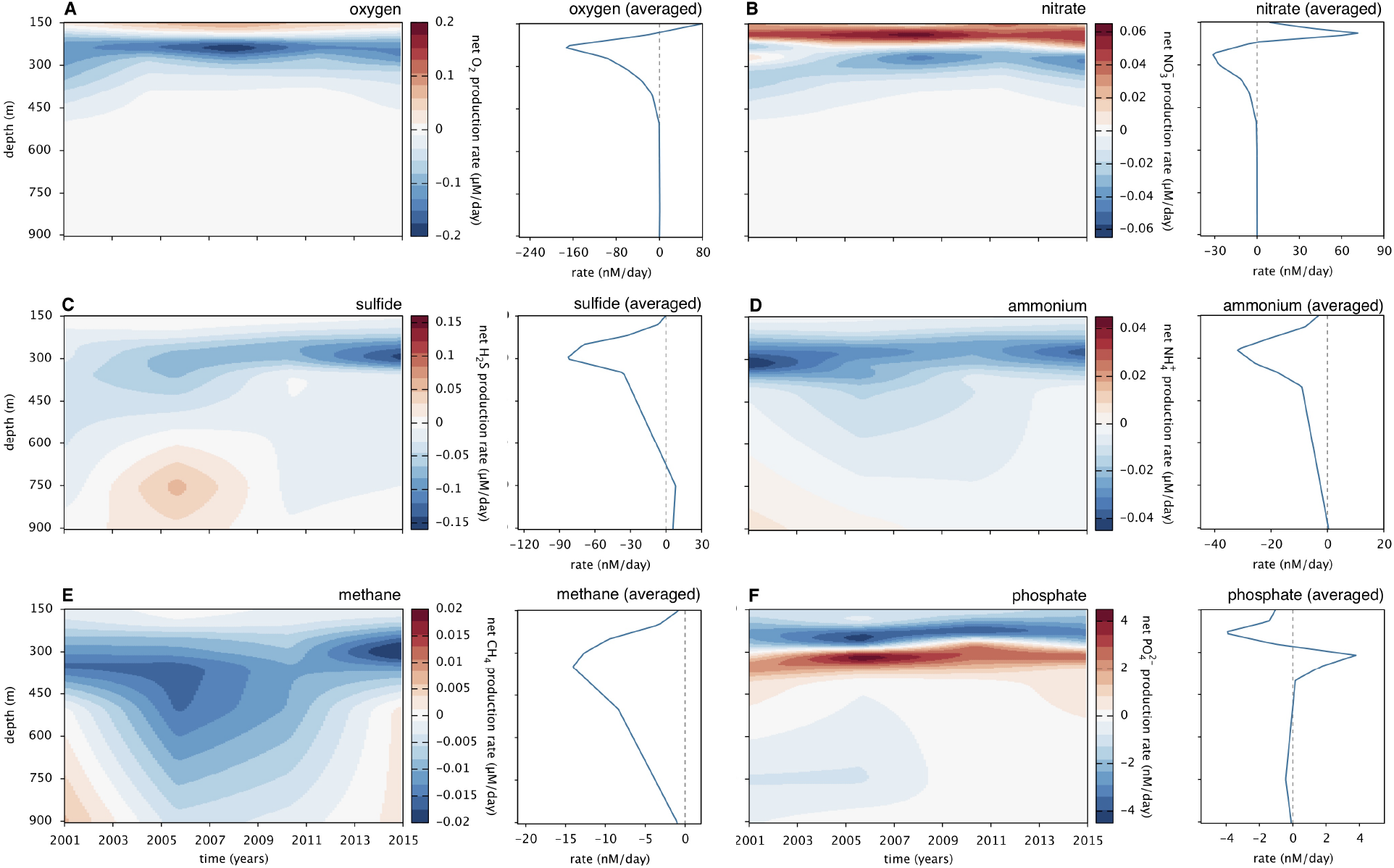
Estimated net metabolite production rates in Cariaco Basin. Volume-specific net metabolite production rates in Cariaco Basin (station CARIACO) over depth and time (contour plots) or averaged over time (depth profiles), estimated via inverse linear transport modeling (A: oxygen, B:nitrate, C:hydrogen sulfide, D:ammonium, E:methane, F:phosphate). In the contour plots, red values correspond to net production, blue values correspond to net consumption, white corresponds to zero net production/consumption. Dashed lines at zero in the time-averaged depth profiles are shown for reference. For estimated gross production and gross consumption rates, see Supplemental Figs. S.8 and S.9, respectively.

Our estimates clearly indicate a production of nitrate near the top (depths 150 m–250 m) and its consumption in the immediately underlying layers (250–300 m, Figs. 3A,B). The weak apparent production of oxygen estimated near the top is likely due to advective (e.g., lateral) transport and/or estimation error, rather than actual *in situ* production at those depths. When integrated across all depths, estimated *in situ* nitrate production near the top almost exactly matches the *in situ* consumption of nitrate immediately below, whereas most of the oxygen consumed in the redoxcline originates from much shallower depths (<150 m, summaries in Table 1). These observations are not surprising, since the main sources of oxygen are the atmosphere and primary production at the surface, while nitrate is likely largely produced by nitrifiers throughout the oxycline wherever ammonium is available and used at depth mostly as an electron acceptor for respiration (Scranton *et al.*, 2014; Cernadas-Martín *et al.*, 2017; Taylor *et al.*, 2018). Using negative values of *R* as an estimate of gross consumption rates, and integrating over all depths, we estimate a gross oxygen consumption of ~13 mmol · m^−2^ · d^−1^ and a gross nitrate consumption of ~2.6 mmol · m^−2^ · d^−1^ on average, indicating that oxygen is a more important terminal electron acceptor in this system than nitrate. Major reductants such as hydrogen sulfide, ammonium and methane diffuse upwards from the bottom layers into the redoxcline where they are largely consumed (Figs. 3C,D,E). The biologically driven flux of electrons from upward diffusing electron donors onto downward diffusing electron acceptors, fuels chemolithoautotrophic microbial activity within the redoxcline (Figs. 4A) and sustains high prokaryotic cell densities (Fig. 4D; Taylor *et al.*, 2006). When integrated (summed) over all considered depths, we estimate an average consumption rate of 11 mmol · m^−2^ · d^−1^ for sulfide, 4.7 mmol · m^−2^ · d^−1^ for ammonium and 3.3 mmol · m^−2^ · d^−1^ for methane, where all area-specific quantities reported here and below are normalized to the basin area at depth 150 m for ease of comparison. The bulk of sulfide, ammonium and methane consumption was found to occur within the redoxcline (overviews in Table 1 and Supplemental Tables S.1 and S.2).

**Table 1:** Estimated mean metabolite fluxes in Cariaco Basin. Estimated *in situ* production and consumption rates, as well as influx and outflux rates across the top (150 m) and bottom boundary (900 m). The total content is depth-integrated, averaged over the considered time interval (years 2001–2014) and measured in mol m^−2^. All rates are depth-integrated where applicable, averaged over the considered time interval, and measured in mmol m^−2^ d^−1^. Depth-integrated or area-specific quantities take into account the variation of the lateral (cross-sectional) basin area with depth, and are normalized to the basin area at depth 150 m to facilitate comparisons. Mean residence times were estimated based on the depth-integrated concentrations and gross input/output rates, using a non-steady-state box model. See Methods for details. For analogous summaries constrained to after 2009, see Supplemental Table S.1. For analogous summaries constrained to the redoxcline (depths 200-400 m), see Supplemental Table S.2.

| variable | $\text{O}_2$ | $\text{NO}_3^{2-}$ | $\text{H}_2\text{S}$ | $\text{NH}_4^+$ | $\text{CH}_4$ | $\text{PO}_4^{3-}$ |
| --- | --- | --- | --- | --- | --- | --- |
| total content (depth-integrated) | 3.5 | 0.81 | 11 | 4.7 | 2.1 | 1.3 |
| gross production rate (depth-integrated) | 2.4 | 2.7 | 1.1 | <0.1 | <0.1 | 0.17 |
| gross consumption rate (depth-integrated) | 13 | 2.7 | 11 | 4.7 | 3.3 | 0.3 |
| net influx rate at top (150 m) | 7.1 | -0.45 | <0.1 | 0.21 | <0.1 | -0.077 |
| gross influx rate at top (150 m) | 7.3 | <0.1 | 0.12 | 0.21 | <0.1 | <0.01 |
| gross outflux rate at top (150 m) | <0.1 | 0.49 | <0.1 | <0.1 | <0.1 | 0.08 |
| net influx rate at bottom (900 m) | <0.1 | <0.1 | 13 | 5.3 | 3.6 | 0.31 |
| gross influx rate at bottom (900 m) | 0.13 | <0.1 | 13 | 5.3 | 3.6 | 0.31 |
| gross outflux rate at bottom (900 m) | 0.17 | <0.1 | <0.1 | <0.1 | <0.1 | <0.01 |
| net influx rate at top+bottom | 7 | -0.43 | 13 | 5.5 | 3.7 | 0.23 |
| total gross influx+production rate | 9.8 | 2.8 | 14 | 5.5 | 3.8 | 0.48 |
| total gross outflux+consumption rate | 14 | 3.2 | 11 | 4.7 | 3.3 | 0.38 |
| mean residence time (years) | 0.67 | 0.69 | 2.6 | 2.7 | 1.7 | 9.1 |

**Figure 4:**
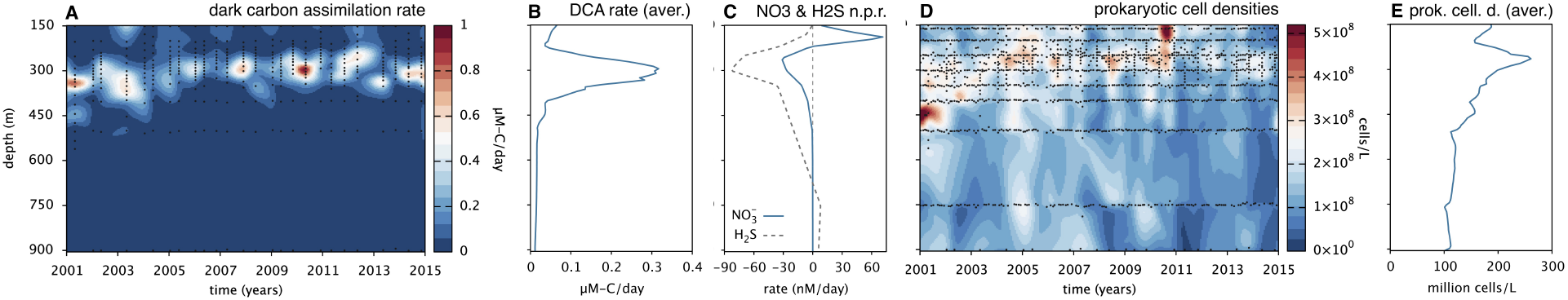
Microbial productivity measured in Cariaco Basin. (A) Dark carbon assimilation (DCA) rate measured in Cariaco Basin (carbon fixed per volume per time) across depth and time. Black dots indicate original data points; the contour plot is obtained via bilinear interpolation. (B) Measured DCA rate, averaged over time (years 2001–2014). (C) Estimated net sulfide and nitrate consumption rates, averaged over time (reproduced from Figs. 3B,C). (D) Measured prokaryotic cell densities (cells per volume) across depth and time. (E) Measured prokaryotic cell densities, averaged over time. Data sources are described in the Methods. For similar figures showing the full water column (including depths <150 m and >900 m) see Supplemental Fig. S.5.

Our estimated sulfide and methane consumption rates are much greater than those estimated by previous studies (~0.1–1.3 mmol · m^−2^ · d^−1^ for sulfide; Taylor *et al.*, 2001; Li *et al.*, 2012b; Taylor *et al.*, 2018 or ~0.04–0.07 mmol · m^−2^ · d^−1^ for methane; Ward *et al.*, 1987; Li *et al.*, 2012b), especially when considering that these previous values would be further reduced after accounting for the smaller basin area at the depths where they were measured (compared to 150 m). This disagreement can be largely explained by the lower diffusivities assumed or estimated in these studies; these studies do acknowledge the great uncertainty in their diffusivity estimates. Our work thus provides a possible explanation for a heavily discussed apparent “conundrum”, whereby sulfide and other electron donor fluxes estimated for Cariaco Basin appeared too low to explain measured dark carbon assimilation (DCA) rates (Taylor *et al.*, 2001; Li *et al.*, 2012b; Jost, 2012; Li *et al.*, 2012a). For example, sulfide fluxes into the redoxcline estimated by Li *et al.* (2012b) are about 50 times lower than ours; assuming a stoichiometric ratio of 1 mol C fixed per mol H_2_S oxidized, Li *et al.* estimated that only 0.2–4.2% of the depth-integrated DCA rate could be explained by sulfide fluxes or, alternatively, that 212 mol-C had to be assimilated per mol-H_2_S oxidized on average. According to our sulfide consumption rate estimate (11 mmol · m^−2^ · d^−1^ on average) and depth-integrated DCA rates (31.6 mmol − C · m^−2^ · d^−1^ on average, Fig. 4A and Taylor *et al.*, 2018), and assuming that sulfide (eventually oxidized to sulfate) is the major source of energy for primary production in the redoxcline (Li *et al.*, 2012b; Taylor *et al.*, 2018), we estimate an average system-wide yield factor of ~2.9 mol C fixed per mol sulfide oxidized. This estimate is still higher than yield factors previously obtained from laboratory cultures of sulfide oxidizers (0.14–0.42 mol C fixed per mol H_2_S; Tuttle and Jannasch, 1979; Kelly, 1990). One explanation may be that energy limitation in Cariaco Basin’s stagnant sub-oxic waters selects for oligotrophic microorganisms, capable of more efficient substrate use than laboratory isolates. Indeed, the energy requirements and efficiencies of various carbon fixation pathways vary widely, depending on the organisms and ecological niches filled (Bar-Even *et al.*, 2011; Berg, 2011; Klatt and Polerecky, 2015). If we follow the thermodynamic arguments by Li *et al.* (2012b), then sulfide-oxidizing chemolithoautotrophic communities in Cariaco Basin may be capable of fixing up to 6.6 mol-C per mol-H_2_S, well above our estimated yield factor. Further, since we ignored the contribution of other electron donors such as ammonium and methane, our estimated yield factor is probably itself an overestimate of the true sulfide-specific yield factor. We also emphasize that this yield factor is an empirical average property of the entire microbial system during the considered time period, and may vary over depth and time depending on environmental conditions and biological interactions. The limited temporal resolution of ILTM-estimated sulfide consumption rates, compared to the rapidly fluctuating measured DCA rates (Fig. 4A), currently hinders a meaningful assessment of the variability of this yield factor.

Most of the sulfide, ammonium and methane input into the system (i.e., via diffusion or *in situ* production) can be attributed to diffusion from the bottom boundary (91%, 95% and 97%, respectively), potentially produced near or in the underlying sediments. Our estimates suggest that some hydrogen sulfide is also produced within the water column (depths ~600–900 m), consistent with the previous detection of sulfate-reducing bacteria in sinking particles at anoxic depths (Suter *et al.*, 2018), although some of the sulfide sources may actually be sulfide diffusing out of the sediments on the basin’s side walls. The contribution of in situ sulfide sources to overall sulfide fluxes into the redoxcline is relatively small (~10%) and has decreased in the latter years, based on the estimated ratio of *in situ* produced versus *in situ* consumed sulfide. A relatively minor contribution of *in situ* sulfide sources is consistent with previous hypotheses (Scranton *et al.*, 1987; Ho *et al.*, 2004). We also found that the majority of phosphate input (gross *in situ* production + influx across the boundaries) is due to diffusion from the bottom boundary (~65%). This phosphate influx from the bottom may partly originate from the remineralization of organic matter in the sediments. Indeed, the estimated ratio of time-averaged ammonium influx vs. phosphate influx from the bottom is ~17:1, closely resembling typical stoichiometric ratios of particulate organic matter in coastal marine ecosystems (17:1 on average; Sterner *et al.*, 2008).

Below the oxic zone, nitrate is presumably used as a terminal electron acceptor by heterotrophic and/or lithotrophic prokaryotes (Scranton *et al.*, 2014; Rodriguez-Mora *et al.*, 2015), fueling complete denitrification to N_2_ (Montes *et al.*, 2013) and/or partial denitrification to intermediates such as nitrite. Since nitrite rarely accumulates below 150 m (Supplemental Fig. S.10), any produced nitrite appears to be re-oxidized to nitrate, further reduced by denitrifiers, or used to anaerobically oxidize ammonium (anammox). The occurrence of denitrification and anammox would be consistent with the reduced ratios of dissolved inorganic nitrogen to phosphorus (N/P) observed in the redoxcline (Muller-Karger *et al.*, 2019), the detection of bacteria capable of various denitrification steps and anammox (Rodriguez-Mora *et al.*, 2015; Cernadas-Martín *et al.*, 2017; Taylor *et al.*, 2018), and similar observations in other oxygen-depleted water columns (Lam and Kuypers, 2011; Lam *et al.*, 2011; Ulloa *et al.*, 2012). Given that sulfide oxidation spatially overlaps substantially with nitrate consumption (Fig. 4C), it is probable that nitrate is at least partly used as a terminal electron acceptor for the oxidation of various sulfur compounds, a process observed in other oxygen-depleted regions of the ocean (Canfield *et al.*, 2010; Schunck *et al.*, 2013; Louca *et al.*, 2016; Rogge *et al.*, 2017). Indeed, the Gammaproteobacterial clades BS-GSO2 and SUP05, members of which are frequently implicated in sulfide oxidation and denitrification in oxygen-poor marine systems (Lavik *et al.*, 2009; Walsh *et al.*, 2009; Fuchsman *et al.*, 2012; Glaubitz *et al.*, 2013; Shah *et al.*, 2017; Rogge *et al.*, 2017), have been observed at high relative abundances in the Cariaco Basin redoxcline (Rodriguez-Mora *et al.*, 2015; Taylor *et al.*, 2018; Suter *et al.*, 2018).

Our estimates reveal a weak but relatively steady consumption of phosphate (0.30 mmol · m^−2^ · d^−1^ on average) between depths ~150–250 m, and a similarly steady production of phosphate (0.17 mmol·m^−2^ ·d^−1^ on average) between depths ~250–350 m (Fig. 3F). This spatially adjacent consumption and production of phosphate leads to the appearance of a subtle phosphate minimum and maximum around the upper and lower half of the redoxcline, respectively. This pattern has been previously partly attributed to a “metal redox shuttle”, whereby phosphate is scavenged during ferrous and manganese oxide formation in the redoxcline and subsequently redissolved at depth (Dellwig *et al.*, 2010; McParland *et al.*, 2015; Muller-Karger *et al.*, 2019). Prokaryotic chemolithoautotrophic activity may also partly drive phosphate consumption within the redoxcline, as suggested by McParland *et al.* (2015) and, in turn, the phosphate production seen immediately below may be due to the remineralization of sinking biomass. The relative importance of such a “biomass shuttle” to the phosphate pool has so far been unclear. Assuming an atomic C:P ratio of 41 for prokaryotic cells (Vrede *et al.*, 2002), and an average dark carbon assimilation rate of 5.25 mmol − C · m^−2^ · d^−1^ between depths 150–250 m (Fig. 4A), one would predict a chemolithoautotrophy-driven phosphate consumption rate of 0.13 mmol · m^−2^ · d^−1^. This prediction is about half of the estimated phosphate consumption rate within that depth range. Hence, a biomass shuttle could only partly explain the phosphate sink and source within the redoxcline, further emphasizing the importance of a putative metal redox shuttle.

We find that the consumption of hydrogen sulfide, methane and, to a lesser extent, ammonium and phosphate has gradually shifted towards shallower depths, and this shift is particularly apparent when comparing times before the year 2010 and afterwards. We also estimate that *in situ* sulfide production at depth substantially decreased over time (Fig. 3C). After 2009, the estimated amount of sulfide produced *in situ* became negligible (<1%) compared to sulfide diffusing from the bottom (summaries in Supplemental Table S.1). Concurrently, sulfide concentrations near the bottom (~1300 m depth) have increased steadily over time (Supplemental Fig. S.11), potentially due to increased production in the underlying sediments, leading to higher diffusive fluxes across the bottom boundary (~10 mmol · m^−2^ · d^−1^ on average before 2010 and ~16 mmol · m^−2^ · d^−1^ afterwards, Supplemental Fig. S.12). This might explain why, despite a decrease of *in situ* sulfide production at depth, net sulfide fluxes into the redoxcline increased (~7.8 mmol · m^−2^ · d^−1^ on average before 2010 and ~9.7 mmol · m^−2^ · d^−1^ afterwards). Interestingly, the upward shift of the redoxcline and the drop of *in situ* sulfide production coincide with major shifts in the composition of the sulfur oxidizing community after 2009 (Taylor *et al.*, 2018). Whether the above changes in nutrient fluxes actually affected, and/or were affected by, changes in the redoxcline-inhabiting community remains unclear.

Three words of caution are warranted. First, due to the limited spatial resolution of our data (and thus, our rate estimates) it is possible that the sinks and sources of metabolites are confined to narrower depth intervals than estimated. Consequently, putatively coupled electron donors (such as sulfide) and electron acceptors (such as oxygen or nitrate), seemingly consumed within the same zone, may in reality be consumed within distinct zones and may only be indirectly coupled through redox shuttles such as manganese and iron (Taylor *et al.*, 2001; Percy *et al.*, 2008). Second, with the data at hand, at each location we can a priori only estimate the local net production rate *R* (gross production minus gross consumption rate), but not the gross production and gross consumption rates separately. It is in principle possible that in some locations some metabolites are both produced and consumed concurrently by separate processes, as observed for sulfate and sulfide in other marine anoxic systems (Canfield *et al.*, 2010). Third, the fact that our estimated rate profiles represent locally averaged net rates implies that, a metabolite produced and consumed in distinct zones but nevertheless in close proximity, may be subject to higher turnover rates than can be inferred from our rate profiles. For example, it is possible that nitrate produced by nitrification is rapidly consumed by denitrification in close proximity immediately below (Cernadas-Martín *et al.*, 2017), and that we thus underestimated nitrate turnover rates in the redoxcline.

### 2.4 Conclusions

We have described a computational approach for estimating vertical fluxes and *in situ* consumption/production rates of dissolved chemical compounds over space and time, via inverse transport modeling. Our approach builds upon established mathematical concepts and has been optimized for water columns or sediments with essentially 1-dimensional geochemical structure, and for which chemical concentrations have been measured at arbitrary spacetime points. We emphasize that despite the apparent simplicity of our models for Cariaco Basin, which assume that eddy diffusion is the dominant mode of salt and metabolite transport in the considered depth interval, our models manage to reproduce the salinity and metabolite concentration data very well (*r*^2^=0.982 for salinity, *r*^2^=0.878–0.973 for metabolites).

We reconstructed vertical fluxes and *in situ* consumption/production rates of several biologically important metabolites in the Cariaco Basin sub-euphotic water column over the course of 14 years. This allowed us to assess the relative importance of *in situ* production in the water column versus supply from (or near) the underlying sediments for various reductants fueling microbial productivity in the redoxcline. By independently estimating the diffusivity in Cariaco Basin over depth and time, rather than relying on parameter values from other marine systems, we further constrained an important source of uncertainty in previous flux estimates (Ho *et al.*, 2004; Li *et al.*, 2012b; Samodurov *et al.*, 2013; Taylor *et al.*, 2018). This revealed that fluxes of various electron donors and acceptors, such as hydrogen sulfide and methane, into the redoxcline are about two orders of magnitude greater than previously estimated (Taylor *et al.*, 2001; Li *et al.*, 2012b; Taylor *et al.*, 2018). Our work thus provides a possible resolution to the long unexplained apparent mismatch between electron donor fluxes and dark carbon assimilation rates in Cariaco Basin (Li *et al.*, 2012b; Jost, 2012; Li *et al.*, 2012a). We also estimated that chemolithoautotrophic activity and remineralization of biomass within the redoxcline only partly explains the phosphate minimum and maximum observed within the redoxcline, thus providing evidence for the existence of an alternative phosphate shuttle. Finally, our work demonstrates that, using appropriate mathematical tools, a wealth of seemingly convoluted information on microbial activity can be extracted from standard chemical concentration time series.

## 3 Methods

### 3.1 Cariaco data

Chemical and physical data from Cariaco station A (coordinates 10.5°N, 64.66°W) were downloaded on April 28, 2018 from the Cariaco Basin time series project website (http://www.imars.usf.edu/cariaco) for station CARIACO. Additional sources of CARIACO chemical data are the NOAA’s National Centers for Environmental Information (NCEI), the Ocean Carbon Data System, the US Biological and Chemical Oceanography Data Management Office (BCO-DMO) and the NASA SeaBASS database. Data collection methods have been described previously (Thunell *et al.*, 2000; Li *et al.*, 2008; Scranton *et al.*, 2014; Muller-Karger *et al.*, 2019). Additional hydrogen sulfide concentration data, recently published by Muller-Karger *et al.* (2019), were obtained directly from the authors. The lateral area of the eastern basin (within which station CARIACO is located) at various discrete depths was taken from Samodurov *et al.* (2013, Table 1), and was linearly interpolated between those depths (Supplemental Fig. S.1).

Inverse linear transport modeling (ILTM) was used to estimate net production rates (*R* in Eq. 6) of oxygen, nitrate, hydrogen sulfide, ammonium, methane and phosphate (dissolved pools). These compounds (referred to here as “metabolites”) were chosen due to their biological importance within the considered depth interval, their relatively good sampling resolution and spatiotemporal coverage, and the fact that their transport across depth can be largely described by eddy diffusion. Nitrite was not included in ILTM because nitrite rarely accumulates to significant levels (Supplemental Fig. S.10); hence, estimated net production rates would be almost zero and dominated by errors despite potentially intense cryptic nitrite fluxes (e.g., as an intermediate of nitrification or denitrification).

### 3.2 Estimating diffusivity in Cariaco Basin

Salinity and temperature profiles were LOESS-smoothened at degree 1 and a span of 10% to reduce noise. Salinity, temperature and pressure profiles were used to calculate the buoyancy frequency (*N*) at each depth and time point, using the R package oce (Kelley, 2014). To reduce noise in the buoyancy frequency stemming from the numerical differentiation of noisy data, the buoyancy frequency was smoothened using a Savitzky-Golay filter of degree 2 along the time axis (Savitzky and Golay, 1964). For any given choice of the parameters *α* and *p* (Eq. 1), we simulated the salinity profile over depth and time by solving the differential equation (2), using the pdepe function in MATLAB^®^. The initial profile was set to the measured salinity profile at the first simulation time point (Jan. 1, 2001). Boundary conditions at the top (150 m) and bottom boundary (900 m) were of Dirichlet type, with the imposed value at each time point and each boundary being the current measured salinity at the boundary’s depth. Salinities between data points were bilinearly interpolated wherever needed for the initial condition and boundary conditions.

We did not account for lateral intrusions of denser, oxygenated water from outside, which are known to occur occasionally in Cariaco Basin (Scranton *et al.*, 2014; Taylor *et al.*, 2018). Salinity and temperature profiles during the time period and depth range considered here do not show obvious signs of foreign water intrusions (Supplemental Figs. S.2A,B). Similarly, oxygen concentration profiles show only weak signs of potential intrusions of oxygenated water at depth (Fig. 2A). It is in principle possible that intrusion events, the subsequent re-equilibration of density structure and the consumption of introduced oxidants all occur at much shorter time scales than resolved by the monthly time series. However, the good agreement of the fitted diffusivity models with the salinity data (*r*^2^ = 0.982, Supplemental Fig. S.3) and the metabolite concentration data (*r*^2^=0.878–0.973) further suggests that lateral water intrusions had little effect on the salt and metabolite budgets within the considered spatiotemporal domain.

The predicted salinity profile *Ŝ* was compared to the measured salinity data (*S*) by means of the fraction of explained variance (*r*^2^), calculated as:

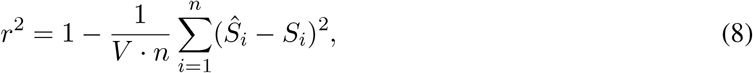

where *i* iterates over all available salinity data points (*n*=111,920), *Ŝ*_*i*_ is the salinity predicted for the same spacetime point as *S*_*i*_ and *V* is the sample variance of the measured salinities *S*_*i*_. The power law parameters *α* and *p* were gradually fitted until *r*^2^ reached a maximum, using the “interior-point” optimization algorithm encoded by the function `fmincon` in MATLAB^®^. To avoid non-global local optima, we repeated the fitting 200 times, each time with randomly chosen initial values for *α* and *p*. The distribution of fitted parameters, as a function of the maximized *r*^2^, is shown in Supplemental Fig. S.13. The parameter set corresponding to the highest *r*^2^ was taken as the final estimate. The same approach was also used to fit the Munk-Anderson diffusivity model (Eq. 3), as well as the combined power-law + Munk-Anderson model (sum of Eqs. 1 and 3). For details on the “anchored” diffusivity estimate (Eq. 4), performed here solely for sanity checking purposes, see Supplement S.1. For the subsequent ILTM analysis, we used the diffusivity obtained from the fitted power-law model.

#### 3.3 Inverse linear transport modeling

Mathematical background and computational details on our ILTM approach are provided in Supplement S.4. Briefly, the differential equation (6) was used to calculate a linear mapping (represented as a matrix 𝕋, see Eq. 7) between any given net production rates (on a finite grid of spacetime points) and the corresponding predicted volumetric concentration profiles (on the same spacetime points as the concentrations measurements). The spatiotemporal grid on which *R* was estimated (“fitting grid”) was chosen separately for each metabolite to account for differences in sampling resolution and spatiotemporal variability of metabolite concentrations, and such that its size was substantially lower than the number of available data points (overview in Supplemental Fig. S.6 and Supplemental Table S.3). In all cases, the number of considered data points was more than 10 times the size of the fitting grid.

Prior to any prediction, the net production rates on the fitting grid were interpolated onto a high-resolution grid (“refined grid”) using an interpolation matrix 𝕀, which maps rates on the fitting grid to rates on the refined grid. Because our numerical differential equation solver only returns solutions on a rectangular spatiotemporal grid (“prediction grid”), an additional interpolation is performed from the prediction grid onto the spacetime points of the data (using a suitable matrix ℙ). Hence, 𝕋 is composed of 3 matrices, 𝕋 = ℙ·𝔾·𝕀, where 𝔾 encodes the “Green’s function” (sometimes called “fundamental solution”) of the differential equation (Duffy, 2001). Each row in the matrix 𝔾 corresponds to the solution of the differential equation (6) evaluated at a specific point on the prediction grid, if the net production rate was zero in all but a single point on the refined grid. The resolutions of the refined grid and the prediction grid were chosen sufficiently high to ensure a high accuracy of the solutions of the differential equation. The estimation of net production rates on the fitting grid can be formulated as a least-squares optimization problem (minimizing the expression in Eq. 7), which we solved using linear algebra routines in MATLAB^®^ (MATLAB, 2010). For each metabolite, the fraction of variance explained by the predicted concentrations (*r*^2^) was calculated as described above for the salinity model.

### 3.4 Estimation of depth-integrated fluxes and area-specific quantities

In all cases, depth-integration of rates and concentrations took into account the variation of the lateral (cross-sectional) basin area with depth (Supplemental Fig. S.1), and all depth-integrated quantities (e.g., production rates) and area-specific quantities (e.g., fluxes through the top and bottom boundaries, or fluxes into the redoxcline) are normalized with respect to the basin area at depth 150 m. For example, if *R*(*t, z*) is the estimated net production rate of some metabolite, then its depth-integrated value within some depth-interval [*z*_1_, *z*_2_], averaged over some time interval [*t*_1_, *t*_2_], was defined as:

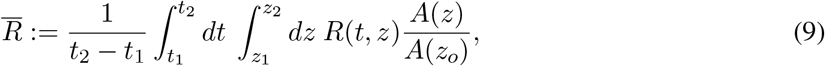

where *A*(*z*) is the lateral basin area and *z*_*o*_=150 m. Thus, in this example, 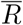 is the hypothetical area-specific vertical flux one would observe at depth *z*_*o*_ if the total number of metabolite molecules produced between depths *z*_1_ and *z*_2_ (integrated over all latitudes/longitudes) was equal to the number of molecules vertically transported past depth *z*_*o*_. The depth *z*_*o*_=150 m was chosen as reference because it is the approximate maximum depth at which Cariaco Basin connects to the ocean, thus marking the Basin’s “upper boundary”, although any other depth could have been used instead.

Net metabolite flux rates across the top (150 m) and bottom boundary (900 m) were estimated from the local metabolite concentration gradients and the local diffusivities. Gross influx rates and gross outflux rates through each boundary were then estimated by using the positive or negative part of the net flux rates, as appropriate. Net *in situ* production rates were estimated via ILTM fitting, as described above. Gross production rates or gross consumption rates were then estimated by taking the positive or negative part of the net production rates, as appropriate. Note that this approach may underestimate actual production and consumption rates, if these occur concurrently at the same depth, since ILTM can a priori only reveal net rates. Estimated *in situ* gross production and gross consumption rates were depth-integrated using the trapezoid rule. A metabolite’s mean total input rate (*R*_*i*_, in mmol · m^−2^ · d^−1^) was defined as the sum of the time-averaged depth-integrated estimated gross production rate plus its time-averaged gross influx rates at the top and bottom boundaries (all normalized to the basin area at depth 150 m). Similarly, a metabolite’s mean total output rate (*R*_*o*_) was defined as the sum of the time-averaged depth-integrated estimated gross consumption rate plus its time-averaged gross outflux rates at the top and bottom boundaries. A metabolite’s total content (*X*, in mol·m^−2^) was calculated by integrating the metabolite’s measured concentration over the entire depth interval while accounting for the variable lateral (cross-sectional) basin area, and subsequently averaged over time. The mean residence time of a metabolite in the considered water column (depths 150–900 m) was estimated from the non-time-averaged total input and output rates (*R*_*i*_(*t*) and *R*_*o*_(*t*)) and the non-time-averaged total content (*X*(*t*)), based on a non-steady-state box model (see Supplement S.3). We mention that, in the case of steady state the mean residence time predicted by the box model would be *X/R*_*o*_; this simplified formula was used previously by Li *et al.* (2012b) under the implicit assumption of steady state. All estimates are listed in Table 1 and Supplemental Tables S.1 and S.2.

## Acknowledgements

S.L. was supported by an NSERC grant and a postdoctoral fellowship from the Biodiversity Research Centre, UBC. M.D. was supported by an NSERC Discovery Grant. We thank all participants and funders of the CARIACO Ocean Time Series program, including the US National Science Foundation for long-term support and in particular through NSF grant OCE-1259110 (to M.I.S. and G.T. Taylor), the Venezuelan Consejo Nacional de Investigaciones Cienti cas y Tecnológicas and the Venezuelan Fondo Nacional de Ciencia y Tecnología. We thank Christopher Wolfe and three anonymous reviewers for comments on our manuscript.

## Author contributions

S.L. conceived the project, wrote the computer code, performed the analyses and wrote a first draft of the manuscript. M.I.S., G.T.T. and Y.M.A. were some of the coordinators of the CARIACO Ocean Time Series program. All authors helped interpret the results, and contributed to the writing of the manuscript.

## Abbreviations

ILTM: inverse linear transport modeling
DCA: dark carbon assimilation

## Data Availability

All raw data used in this article have been published previously (Scranton *et al.*, 2014; Muller-Karger *et al.*, 2019) and are publicly available at the Cariaco Basin Time Series project website (http://www.imars.usf.edu/cariaco).

## Code availability

Our MATLAB^®^ code for estimating diffusivity based on salinity profiles and for estimating metabolite fluxes via ILTM is available online at: www.loucalab.com/archive/CariacoMetabolic

## Competing financial interests

The authors declare that they have no competing interests.

